# Parasitism does not reduce thermal limits in the intermediate host of a bopyrid isopod

**DOI:** 10.1101/2023.03.31.535176

**Authors:** Matthew Sasaki, Charles Woods, Hans G. Dam

## Abstract

Parasitism has strong effects on community dynamics. Given the detrimental effects parasites have on host health, infection or infestation might be expected to reduce upper thermal limits, increasing the vulnerability of host species to future climate change. Copepods are integral components of aquatic food webs and biogeochemical cycles. They also serve as intermediate hosts in the life cycle of parasitic isopods in the family Bopyridae. Given the important effects both copepods and isopod parasites play in aquatic communities, it is important to understand how the interaction between parasite and host affects thermal limits in order to better predict how community dynamics may change in a warming climate. Here we examined the effect of infestation by larvae of a bopyrid isopod on cosmopolitan copepod *Acartia tonsa* to test the hypothesis that infestation reduces thermal limits. To aid with this work, we developed an affordable, highly portable system for measuring critical thermal maxima of small ectotherms. We also used meta-analysis to summarize the effects of parasitism on critical thermal maxima in a wider range of taxa to help contextualize our findings. Contrary to both our hypothesis and the results of previous studies, we observed no reduction of thermal limits by parasitism in *A. tonsa*. These results suggest that life history of the host and parasite may interact to determine how parasite infestation affects environmental sensitivity.

## Introduction

Ecological dynamics are shaped by species interactions, which are in turn affected by the environmental context. Understanding environmental sensitivities can therefore provide important insights into the responses of communities to climate change. Species interactions can, however, also exert an influence on organismal sensitivity to environmental conditions. Parasitism, for example, can have a wide range of effects on community dynamics (Hatcher et al., 2012; Thomas et al., 2005). By nature of its detrimental effects on host health and energy reserves, parasitism might also be expected to affect ecological dynamics by modifying host sensitivity to changing conditions (Hector et al., 2021). In particular, vulnerability to temperature extremes is important to understand given the rapid increases in average temperatures and the increase in the frequency and intensity of extreme events like heatwaves (Frölicher and Laufkötter, 2018; Johnson and Lyman, 2020).

For a limited range of host and parasite taxa, previous studies have observed a decreased host capacity to tolerate increased temperatures as a result of parasitization (Hector et al., 2021). More generally, however, parasites have been shown to elicit a wide range of molecular, physiological, behavioral, and life history responses in the host (Burris and Dam, 2014; Díaz-Morales et al., 2023; Doublet et al., 2017; Frank et al., 2013; Fuess et al., 2021; Heil, 2016; Hurd, 2001; Laverty et al., 2017; Wertheim et al., 2005; Wertheim, 2022). These diverse responses may alter how parasitism affects host thermal limits. For example, parasitization-induced increases in heat shock protein expression may enable the host to better respond to acute temperature stress (Encomio and Chu, 2007; Selbach et al., 2020). As parasitism is widely observed across taxa, examining the diverse responses to infection or infestation and how these feedback onto environmental sensitivity may have important consequences for our understanding of how communities will respond to climate change.

Copepods are the most abundant pelagic metazoans in the ocean, and play important ecological and biogeochemical roles in coastal marine systems. Pelagic calanoid copepods also serve as the intermediate host for parasitic isopods in the family Bopyridae (Mauchline, 1998), some of which infest commercially harvested decapods as their final host (Conner and Bauer, 2010; Roccatagliata and Lovrich, 1999; Shields, 2012; Somers and Kirkwood, 1991; Vinuesa and Balzi, 2010). As in other systems, less attention has been paid to the effects these parasites have on their intermediate hosts. *Acartia tonsa* (Dana, 1849) is foundational species across the coastal Western Atlantic (Turner, 1981), and is known to be the intermediate host for several species of bopyrid isopods (Dale and Anderson, 1982; Williams et al., 2022). Previous work has shown that parasitism by isopod larvae reduces metabolic rates in A. *tonsa* (Anderson, 1975). In other species of copepods, parasitism by isopod larvae induces infertility (Uye and Murase, 1997). While it is clear that infestation induces a physiological response, it is unknown whether parasitism also affects the temperature sensitivity of these copepod hosts. In rapidly warming coastal systems, parasitism-induced reductions in host thermal limits may have substantial effects on food web and biogeochemical dynamics.

Given that *Acartia tonsa* is a foundational species, we aim to test the hypothesis that infestation by bopyrid isopod larvae reduces its thermal limits. We also contextualize our findings using a meta-analysis of the effects of parasitism on critical thermal maxima. Surprisingly, we find that bopyrid infestation does not reduce thermal limits in *A. tonsa* females. This is in stark contrast to previous studies, that generally report parasitism reduces critical thermal maxima. This may reflect differential effects of parasites on the temperature sensitivity of final and intermediate hosts. In any case, the deviation of our results from those of previous studies highlights the importance of examining the effects of parasitism on thermal limits across diverse taxa.

## Methods

### Animal Collection

Copepods were collected from Key Largo, Florida in late February 2023 (25.283775N, −80.330165W; water temperature: 27°C; salinity: 27 psu). A substantial portion of adult *A. tonsa* females bore larvae of a bopyrid isopod attached to their prosome (Figure 1), likely *Probopyrus floridensis* (Dale and Anderson, 1982). While not the initial target of this sampling effort, this allowed us to opportunistically examine the effects of parasitism on individual thermal limits. After collection, the contents of the plankton tow were maintained at 22°C. An aquarium bubbler was used to ensure constant aeration. Mature *Acartia tonsa* females were sorted from the bulk contents of the plankton tow, with those females bearing a bopyrid isopod larva attached to their prosome kept separate. No more than one isopod was present per female. Individual thermal limits were measured in batches of ten (five each with and without isopods).

**Figure 1:**
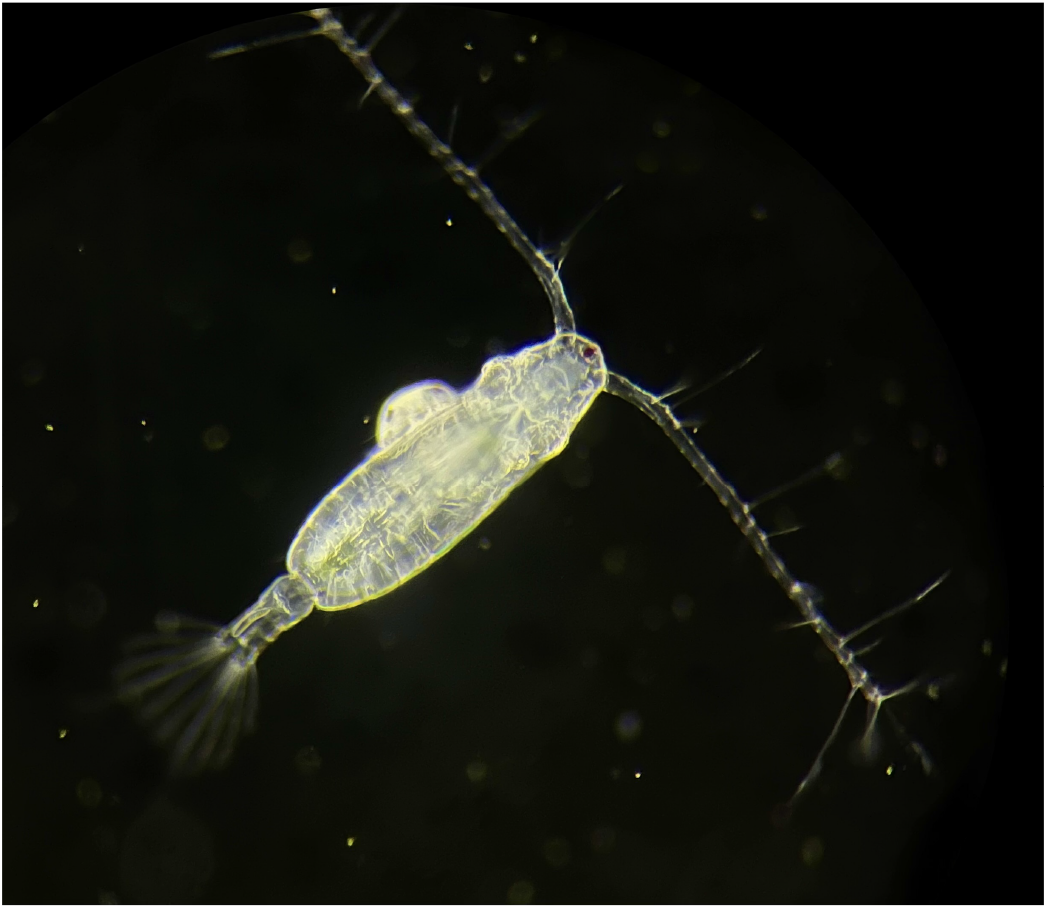
A mature Acartia tonsa female bearing a bopyrid isopod larva. Image was taken after returning from the field site using an Olympus stereomicroscope. Photo credit: Matthew Sasaki

### CT_max_ Device Description

Critical thermal maxima (CT_max_) are measured using a dynamic ramping assay, and indicate the maximum temperature at which an individual can maintain normal activity (Cowles and Bogert, 1944; Lutterschmidt and Hutchison, 1997). These measurements are commonly used across a wide range of taxa (Bennett et al., 2018). Copepod thermal limit measurements are still relatively uncommon, however, despite their high abundance, widespread distribution, and ecological importance (Sasaki and Dam, 2021).

We have developed a highly portable system to assist with making thermal limit measurements in copepods and other small ectotherms, suitable for both rapid measurements under field conditions or controlled laboratory settings. The system has three components: a reservoir, a water bath, and a temperature sensor (Figure 2). We used a 5-gallon bucket covered with a neoprene sleeve as our reservoir. The reservoir is filled with ~15 L of water, which is slowly warmed using a 300-watt aquarium water heater. Temperature ramping rate is determined by the interaction between the power output of the aquarium heater and the volume of water in the reservoir, enabling users to alter ramping rates to suit their needs. The reservoir also contains two aquarium pumps, one of which vigorously circulates water within the reservoir while the other pumps water from the reservoir into the water bath, which sits atop the reservoir. The water bath is a transparent plexiglass box that fits over the opening of the reservoir. Water is pumped up from the reservoir, flooding the water bath. A recession cut into one edge allows water to spill back into the reservoir. The water bath contains several test tube racks that are used to secure the experimental vessels (50 mL flat-bottom glass vials). Because the box is transparent, individuals are easily monitored through the side of the water bath throughout the trial, eliminating the necessity to remove experimental vessels and any resulting temperature fluctuations. A small Arduino computer system logs temperature with three independent sensors at 5 second intervals. These sensors are small enough to be placed inside the glass vials, providing a continuous record of the temperatures throughout the assay. The data are stored on a microUSB card for easy access. All required materials, code, and technical specifications are maintained in a separate data repository (https://github.com/ZoopEcoEvo/CTmax_device). A major advantage of this system is the portability - most of the components fit within the empty five-gallon bucket, allowing for secure transport or shipping. The cost of the unit is also relatively low.

**Figure 2:**
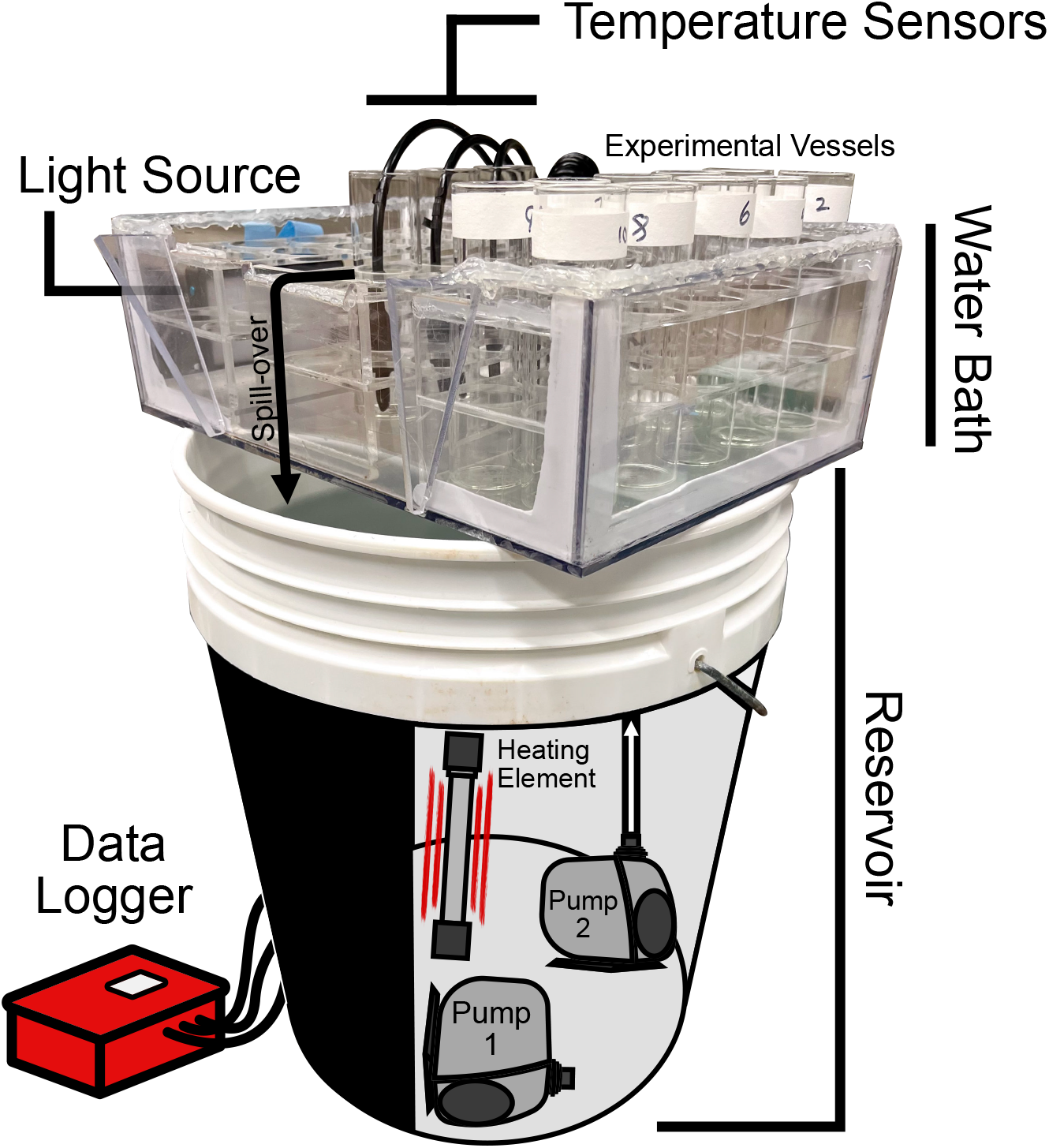
The different components of the setup used to measure CTmax values. The reservoir contains two water pumps and a 300-watt fixed output heating element. Pump 1 is placed parallel to the base of the reservoir and vigorously circulates the water. Pump 2 is used to move water from the reservoir up into the water bath. As the bath is flooded, water spills back over into the reservoir through a cutout in the front face. The water bath contains several test tube racks which hold the experimental vessels (50 mL flat-bottom glass tubes) in place. Several other of these tubes are placed throughout the bath to hold the temperature sensors. Temperature is recorded by these three sensors to a data logger, run by a small Arduino computer system. A small light source is attached across from the experimental vessels to provide backlighting for the experimental vessels, assisting with observation during the experiment.

### Measuring CT_max_

We selected a ramping rate of ~0.2°C per minute. Similar ramping rates have been used previously to estimate CT_max_ in copepods (Harada and Burton, 2019; Jiang et al., 2009). To initiate each CT_max_ assay, ten copepods (five bearing isopods and five without) were placed individually into glass tubes with 10 mL of 0.2 um filtered seawater, collected from the same location as the copepods and pre-acclimated to the same temperature copepods were maintained at. After a brief resting phase, the water heater was turned on, initiating the temperature ramp. Simultaneously, the temperature logger began to record temperature and a stopwatch began recording the time elapsed. Individuals were then continuously monitored as water temperature increased. Copepods not actively moving were checked by rotating the glass vial, causing slight water movement. In active copepods, this stimulus is sufficient to elicit a jump response. The time at which an individual stopped responding to this stimulus was recorded to the nearest second. In addition to unresponsiveness, other indications that an individual had reached their CT_max_ were i) a sustained position on the floor of the vessel, ii) antennules that had curved inwards, were held flat against the prosome, or were held at abnormal backwards angles, and iii) a backwards arching of the urosome above the prosome. After an individual reached its CT_max_, the glass vial it was held in was removed from the water bath.

Once the assay was completed, the recorded times were converted to individual CT_max_ values in °C using the continuous temperature record logged by the Arduino system. We used this time-based method instead of directly monitoring the temperature for two reasons: 1) it was faster to record the time an individual stopped responding than to check the temperature readings from the three sensors, increasing the resolution of our measurements; and 2) it reduced sub-conscious bias stemming from past knowledge or expectations about copepod thermal limits. To estimate CT_max_ we averaged the temperature readings from the three individual sensors over a period directly preceding the time the individual ceased responding to stimuli. This period extended from the time at which an individual was recorded as having stopped responding to stimulus to the last time the individual was checked. As it takes around 5 seconds to check each individual for a response, the duration of this window was estimated for each individual as the number of vials remaining in the water bath when it reached its CT_max_, multiplied by 5 seconds. Thus, the uncertainty window decreased in length as the trial went on, until for the final individual the window included just the amount of time it took to check whether the individual had stopped responding.

### Statistical Analysis

All statistical analyses were performed with R (Version 4.1.3; R Core Team 2022). The CT_max_ values for the two groups (infested and non-infested) were compared using mean difference as an effect size estimate. 95% confidence intervals were estimated using non-parametric bootstrapping (Ho et al., 2019). Since multiple replicate experiments were run and CT_max_ may have changed as time since collection increased, we also ran a linear mixed effects model, with CT_max_ modeled as a function of infection status with experimental replicate as a random effect.

### Meta-Analysis

In order to compare the effects we observed here with previous work, we also conducted a meta-analysis. Potential studies were obtained with a Web of Science search for “(CTmax OR”upper thermal limit” OR “thermal tolerance”) AND parasit*” on March 6th, 2023. This yielded 132 results, which were individually screened for inclusion in the meta-analysis. Given the small number of studies returned by the search, we examined the entire text when determining study inclusion. In order to be included, a study must have reported CTmax values for both infected and uninfected host animals, along with sample sizes and standard deviations or errors. We used strict inclusion criteria to maximize comparability (excluding studies that measured thermal limits in time to knockdown, or percent survivorship after static temperature exposure, etc.). A total of five previously published studies met these criteria. In cases were studies reported multiple experimental treatments (e.g. ramping rates) only one set of measurements was included per host-parasite pair, selected to represent intermediate conditions. Mean thermal limits, standard deviations, and sample sizes were extracted from the study text or tables, although wherever possible these values were calculated using the raw data. We then estimated standardized mean difference and 95% confidence interval for each comparison between infected and non-infected animals using the metafor package (Viechtbauer, 2010).

## Results

### CT_max_ Values

Using the set up described above, we were able to gradually increase the temperature of the water within the vials (Figure 3a). Ramping rates are always within a 0.1-0.3°C per minute range, but due to the imperfect insulation of the reservoir, these rates decreased slightly over time from ~0.25°C to ~0.2°C (Figure 3b). Contrary to expectations, we did not observe a decrease in thermal limits in the infested copepods (Figure 4). Instead, the observed effect of bopyrid infestation on CT_max_ was slightly positive, with a confidence interval that strongly overlapped zero (mean difference: 0.1 °C; 95% CI −0.21 °C to 0.35°C). The linear mixed effect model also indicated no effect of bopyrid infestation (p = 0.48). This result is strongly affected by three parasitized individuals with substantially lower thermal limits (Figure 4). When these thermal limits are removed, there is a small positive effect of infestation (mean effect: 0.3°C; 95% CI 0.098°C to 0.51 °C). With the three low values removed, the linear mixed effects model also indicates a positive effect of bopyrid infestation on CT_max_ (p = 0.006).

**Figure 3:**
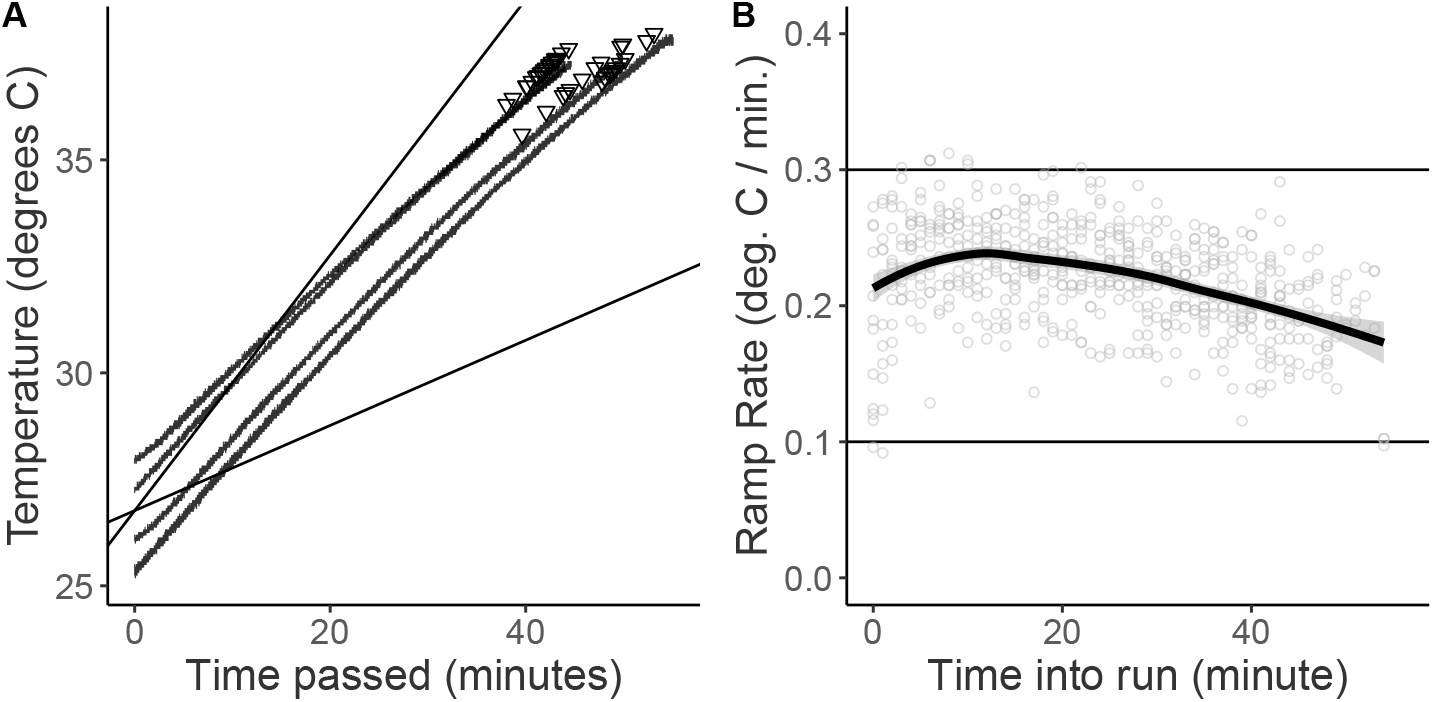
Temperature data from each of the four CTmax assays. A) The average temperature recorded across the three sensors over time. Triangular points indicate the points at which individuals were observed to reach their critical thermal maximum. B) The ramping rate observed during the CTmax assays. Temperature data were binned to minute intervals. The ramping rate was then calculated as the linear change over time using all measurements recorded by the three sensors within that minute interval.

**Figure 4:**
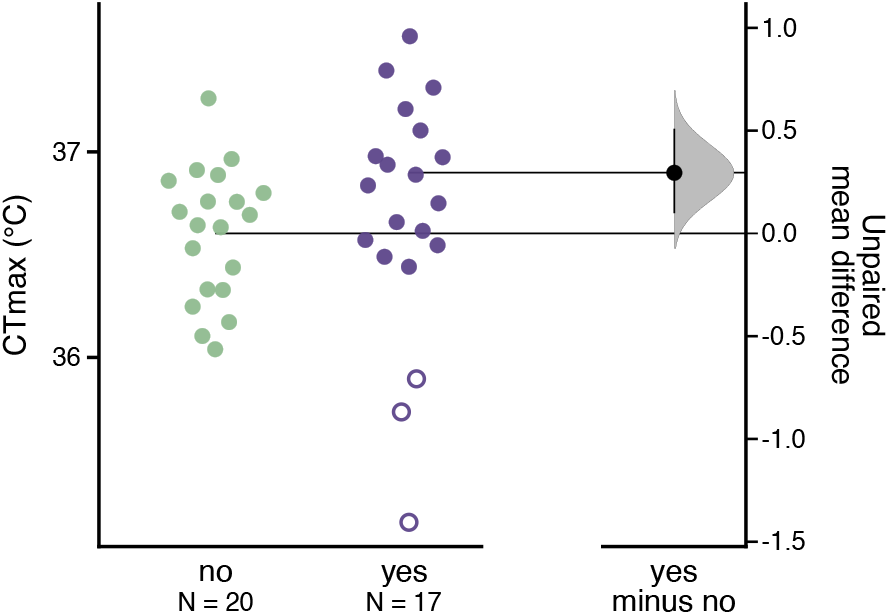
Estimation plot showing observed CTmax values and the estimated effect of infestation. Infestation status is represented using different colored points (no bopyrid in green; with a bopyrid (yes) in purple). The three low thermal limits that were excluded are shown as unfilled symbols within the infested group, and are not included in the sample sizes.

### Meta-Analysis

Five studies met our criteria for inclusion in the meta-analysis, yielding 15 contrasts between parasitized and unparasitized host animals. The studies included in the meta-analysis generally focused on amphibian parasitization by fungal or protistan parasites (Fernández-Loras et al., 2019; Greenspan et al., 2017; Sherman, 2008), but also included studies examining an arthropod host and bacterial parasite (Hector et al., 2019), and, as examined in our study, an arthropod host with an arthropod parasite (Agosta et al., 2018). These studies generally reported a negative effect of parasitism on CT_max_, although in several cases confidence intervals overlapped zero (Figure 5). Note that each of the five previous studies examined the effect of parasites on their final host, while *A. tonsa* is the intermediate host for this isopod.

**Figure 5:**
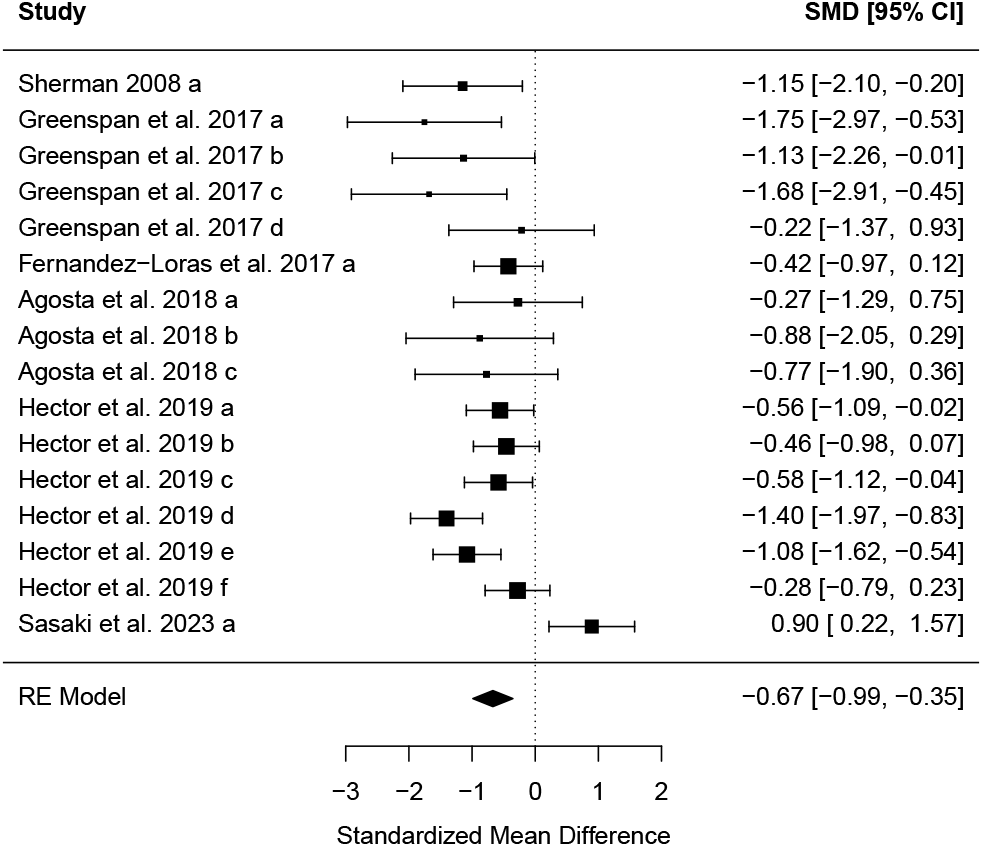
A forest plot showing the standardized mean difference estimate for each comparison of CTmax in infected and non-infected hosts, along with the 95 percent confidence interval. Multiple contrasts from the same study are indicated by different letters following the study ID. The size of each point is proportional to the weight of each contrast. The diamond at the bottom of the plot summarizes the overall effect estimate.

## Discussion

We observed that, contrary to our hypothesis, bopyrid infestation does not have a strong negative effect on the thermal limits of *Acartia tonsa* females. These findings were also contrary to the results of previously published studies on the effects of parasitism on the critical thermal limits of a host.

While our results clearly refute that parasitism decreases thermal limits in *Acartia tonsa*, the observation of a positive effect is strongly affected by whether we include or exclude three notably lower thermal limit measurements in the infested treatment. Assuming individuals are not switching hosts, females bearing larger bopyrids are likely older than females with smaller bopyrids. Visual inspection before the beginning of the experiment indicated that the three individuals with particularly low thermal limits all hosted larger bopyrids; the low thermal limits may therefore reflect the older age of these females. As we were opportunistically measuring the effects of bopyrids, we lacked the required equipment to measure individual body lengths after the CTmax measurements were made and are unable to test for this correlation between larval length and host CTmax. It is also possible that the effect of infestation increases with parasite size or developmental stage (Pike pers. comm. in Marshall and Orr, 1972), and that isopod parasitism has stage-specific effects on host thermal limits. These dynamics are worth further examination in more targeted studies.

Also worth examining are the relative effects of bopyrid infestation on the thermal limits of other copepod species. While *Acartia tonsa* commonly serves as the intermediate host, other species have also been observed to be infested by bopyrid larvae (Owens and Rothlisberg, 1995; Pike, 1961; Uye and Murase, 1997; Williams et al., 2022). Copepods are also subject to infestation or infection by a diverse range of other taxa (Bass et al., 2021). Examining the effects of these other taxa on copepod thermal limits is also crucial. Both variation in the effects of parasitism across species (Hatcher et al., 2006, 2008) and differences in thermal sensitivity (Willett, 2010) can have strong effects on community dynamics by reducing or intensifying competitive interactions. Increased relative susceptibility of other species to isopod parasitism, for example, may further promote the dominance of *Acartia tonsa* in these coastal communities.

The lack of a strong negative effect of parasitism on CT_max_ in *Acartia tonsa* was contrary to our hypothesis that infection would reduce thermal limits. It is possible that infestation stimulated the production of heat shock proteins (Encomio and Chu, 2007; Frank et al., 2013; Yang et al., 2022; Yu et al., 2021), a key component of the copepod heat stress response (Rahlff et al., 2017; Schoville et al., 2012). We wonder, however, if perhaps our initial hypothesis should have instead been that parasitism does not reduce the thermal limits of an intermediate host species. Parasites often have complex life cycles, relying on one or more intermediate hosts before locating their final host. In a variable thermal environment, reduction of the host thermal limits by infection or infestation could disrupt the parasite’s development if exposure to high temperature results in host mortality. It may be beneficial, therefore, for parasites to manipulate host physiology in such a way to make these disruptions less likely (Lefèvre et al., 2008). Aphids serving as a vector for a plant virus (analogous to an intermediate host), for example, had increased thermal limits, allowing them to exploit warmer microhabitats than competitors, ultimately aiding in the spread of the virus (Porras et al., 2020). In animal parasites, shifts in host preference toward warmer temperatures (Bates et al., 2011) or more exposed, illuminated sites (MacNeil et al., 2003) may also contribute to shifts in temperature tolerance via acclimation effects. Given the necessity of a pelagic intermediate host for the successful development of this isopod, from an evolutionary perspective, it may make sense that infestation would not reduce host thermal limits.

## Acknowledgements

We thank Ewaldo Leitao and Dail Laughinghouse for providing assistance and laboratory space while collecting animals and running experiments. This work was supported by the National Science Foundation [award numbers 1947965 and 2205848] and a Postdoctoral Seed Award from the University of Connecticut.

## Notes

https://github.com/ZoopEcoEvo/tonsa_infected_CTmax

### Competing Interest Statement

The authors have declared no competing interest.

## References

Agosta, S.J., Joshi, K.A., Kester, K.M., 2018. Upper thermal limits differ among and within component species in a tritrophic host-parasitoid-hyperparasitoid system. PLOS ONE 13, e0198803. http://doi.org/10.1371/journal.pone.0198803.

Anderson, G., 1975. Larval metabolism of the epicaridian isopod parasite probopyrus pandalicola and metabolic effects of p. pandalicola on its copepod intermediate host acartia tonsa. Comparative Biochemistry and Physiology Part A: Physiology 50, 747–751. http://doi.org/10.1016/0300-9629(75)90140-1.

Bass, D., Rueckert, S., Stern, R., Cleary, A.C., Taylor, J.D., Ward, G.M., Huys, R., 2021. Parasites, pathogens, and other symbionts of copepods. Trends in Parasitology 37, 875–889. http://doi.org/10.1016/j.pt.2021.05.006.

Bates, A.E., Leiterer, F., Wiedeback, M.L., Poulin, R., 2011. Parasitized snails take the heat: a case of host manipulation? Oecologia 167, 613–621. http://doi.org/10.1007/s00442-011-2014-0.

Bennett, J.M., Calosi, P., Clusella-Trullas, S., Martínez, B., Sunday, J., Algar, A.C., Araújo, M.B., Hawkins, B.A., Keith, S., Kühn, I., Rahbek, C., Rodríguez, L., Singer, A., Villalobos, F., Ángel Olalla-Tárraga, M., Morales-Castilla, I., 2018. Globtherm, a global database on thermal tolerances for aquatic and terrestrial organisms. Scientific Data 5, 180022. http://doi.org/10.1038/sdata.2018.22.

Burris, Z.P., Dam, H.G., 2014. Deleterious effects of the ciliate epibiont zoothamnium sp. on fitness of the copepod acartia tonsa. Journal of Plankton Research 36, 788–799. http://doi.org/10.1093/plankt/fbt137.

Conner, S.L., Bauer, R.T., 2010. Infection of adult migratory river shrimps, macrobrachium ohione, by the branchial bopyrid isopod probopyrus pandalicola: Parasitism in amphidromous shrimps. Invertebrate Biology 129, 344–352. http://doi.org/10.1111/j.1744-7410.2010.00210.x.

Cowles, R.B., Bogert, C.M., 1944. A preliminary study of the thermal requirements of desert reptiles. Bulletin of the American Museum of Natural History 83, 261–296.

Dale, W.E., Anderson, G., 1982. Comparison of morphologies of probopyrus bithynis, p. floridensis, and p. pandalicola larvae reared in culture (isopoda, epicaridea). Journal of Crustacean Biology 2, 392–409. http://doi.org/10.2307/1548055.

Díaz-Morales, D.M., Bommarito, C., Knol, J., Grabner, D.S., Noè, S., Rilov, G., Wahl, M., Guy-Haim, T., Sures, B., 2023. Parasitism enhances gastropod feeding on invasive and native algae while altering essential energy reserves for organismal homeostasis upon warming. Science of The Total Environment 863, 160727. http://doi.org/10.1016/j.scitotenv.2022.160727.

Doublet, V., Poeschl, Y., Gogol-Döring, A., Alaux, C., Annoscia, D., Aurori, C., Barribeau, S.M., Bedoya-Reina, O.C., Brown, M.J.F., Bull, J.C., Flenniken, M.L., Galbraith, D.A., Genersch, E., Gisder, S., Grosse, I., Holt, H.L., Hultmark, D., Lattorff, H.M.G., Le Conte, Y., Manfredini, F., McMahon, D.P., Moritz, R.F.A., Nazzi, F., Niño, E.L., Nowick, K., van Rij, R.P., Paxton, R.J., Grozinger, C.M., 2017. Unity in defence: honeybee workers exhibit conserved molecular responses to diverse pathogens. BMC Genomics 18. http://doi.org/10.1186/s12864-017-3597-6.

Encomio, V., Chu, F., 2007. Heat shock protein (hsp70) expression and thermal tolerance in sublethally heat-shocked eastern oysters crassostrea virginica infected with the parasite perkinsus marinus. Diseases of Aquatic Organisms 76, 251–260. http://doi.org/10.3354/dao076251.

Fernández-Loras, A., Boyero, L., Correa-Araneda, F., Tejedo, M., Hettyey, A., Bosch, J., 2019. Infection with batrachochytrium dendrobatidis lowers heat tolerance of tadpole hosts and cannot be cleared by brief exposure to ctmax. PLOS ONE 14, e0216090. http://doi.org/10.1371/journal.pone.0216090.

Frank, S.N., Godehardt, S., Nachev, M., Trubiroha, A., Kloas, W., Sures, B., 2013. Influence of the cestode ligula intestinalis and the acanthocephalan polymorphus minutus on levels of heat shock proteins (hsp70) and metallothioneins in their fish and crustacean intermediate hosts. Environmental Pollution 180, 173–179. http://doi.org/10.1016/j.envpol.2013.05.014.

Frölicher, T.L., Laufkötter, C., 2018. Emerging risks from marine heat waves. Nature Communications 9, 650. http://doi.org/10.1038/s41467-018-03163-6.

Fuess, L.E., Weber, J.N., den Haan, S., Steinel, N.C., Shim, K.C., Bolnick, D.I., 2021. Between-population differences in constitutive and infection-induced gene expression in threespine stickleback. Molecular Ecology 30, 6791–6805. http://doi.org/10.1111/mec.16197.

Greenspan, S.E., Bower, D.S., Roznik, E.A., Pike, D.A., Marantelli, G., Alford, R.A., Schwarzkopf, L., Scheffers, B.R., 2017. Infection increases vulnerability to climate change via effects on host thermal tolerance. Scientific Reports 7, 9349. http://doi.org/10.1038/s41598-017-09950-3.

Harada, A.E., Burton, R.S., 2019. Ecologically relevant temperature ramping rates enhance the protective heat shock response in an intertidal ectotherm. Physiological and Biochemical Zoology 92, 152–162. http://doi.org/10.1086/702339.

Hatcher, M.J., Dick, J.T., Dunn, A.M., 2008. A keystone effect for parasites in intraguild predation? Biology Letters 4, 534–537. http://doi.org/10.1098/rsbl.2008.0178.

Hatcher, M.J., Dick, J.T., Dunn, A.M., 2012. Diverse effects of parasites in ecosystems: linking interdependent processes. Frontiers in Ecology and the Environment 10, 186–194. http://doi.org/10.1890/110016.

Hatcher, M.J., Dick, J.T.A., Dunn, A.M., 2006. How parasites affect interactions between competitors and predators. Ecology Letters 9, 1253–1271. http://doi.org/10.1111/j.1461-0248.2006.00964.x.

Hector, T.E., Sgrò, C.M., Hall, M.D., 2019. Pathogen exposure disrupts an organism’s ability to cope with thermal stress. Global Change Biology 25, 3893–3905. http://doi.org/10.1111/gcb.14713.

Hector, T.E., Sgrò, C.M., Hall, M.D., 2021. Thermal limits in the face of infectious disease: How important are pathogens? Global Change Biology 27, 4469–4480. http://doi.org/10.1111/gcb.15761.

Heil, M., 2016. Host manipulation by parasites: Cases, patterns, and remaining doubts. Frontiers in Ecology and Evolution 4. http://doi.org/10.3389/fevo.2016.00080.

Ho, J., Tumkaya, T., Aryal, S., Choi, H., Claridge-Chang, A., 2019. Moving beyond p values: data analysis with estimation graphics. Nature Methods 16, 565–566. http://doi.org/10.1038/s41592-019-0470-3.

Hurd, H., 2001. Host fecundity reduction: a strategy for damage limitation? Trends in Parasitology 17, 363–368. http://doi.org/10.1016/s1471-4922(01)01927-4.

Jiang, Z.B., Zeng, J.N., Chen, Q.Z., Huang, Y.J., Liao, Y.B., Xu, X.Q., Zheng, P., 2009. Potential impact of rising seawater temperature on copepods due to coastal power plants in subtropical areas. Journal of Experimental Marine Biology and Ecology 368, 196–201. http://doi.org/10.1016/j.jembe.2008.10.016.

Johnson, G.C., Lyman, J.M., 2020. Warming trends increasingly dominate global ocean. Nature Climate Change 10, 757–761. http://doi.org/10.1038/s41558-020-0822-0.

Laverty, C., Brenner, D., McIlwaine, C., Lennon, J.J., Dick, J.T., Lucy, F.E., Christian, K.A., 2017. Temperature rise and parasitic infection interact to increase the impact of an invasive species. International Journal for Parasitology 47, 291–296. http://doi.org/10.1016/j.ijpara.2016.12.004.

Lefèvre, T., Roche, B., Poulin, R., Hurd, H., Renaud, F., Thomas, F., 2008. Exploiting host compensatory responses: the ‘must’ of manipulation? Trends in Parasitology 24, 435–439. http://doi.org/10.1016/j.pt.2008.06.006.

Lutterschmidt, W.I., Hutchison, V.H., 1997. The critical thermal maximum: history and critique. Canadian Journal of Zoology 75, 1561–1574. http://doi.org/10.1139/z97-783.

MacNeil, C., Fielding, N.J., Hume, K.D., Dick, J.T., Elwood, R.W., Hatcher, M.J., Dunn, A.M., 2003. Parasite altered micro-distribution of gammarus pulex (crustacea: Amphipoda). International Journal for Parasitology 33, 57–64. http://doi.org/10.1016/s0020-7519(02)00229-1.

Marshall, S.M., Orr, A.P., 1972. The Biology of a Marine Copepod: Calanus finmarchicus (Gunnerus). Springer Berlin Heidelberg, Berlin, Heidelberg.

Mauchline, J., 1998. The Biology of Calanoid Copepods. volume 33 of Advances in Marine Biology. Academic Press, San Diego.

Owens, L., Rothlisberg, P., 1995. Epidemiology of cryptonisci (bopyridae:isopoda) in the gulf of carpentaria, australia. Marine Ecology Progress Series 122, 159–164. http://doi.org/10.3354/meps122159.

Pike, R.B., 1961. Observations on epicaridea obtained from hermit-crabs in british waters, with notes on the longevity of the host-species. Annals and Magazine of Natural History 4, 225–240. http://doi.org/10.1080/00222936108651101.

Porras, M.F., Navas, C.A., Marden, J.H., Mescher, M.C., De Moraes, C.M., Pincebourde, S., Sandoval-Mojica, A., Raygoza-Garay, J.A., Holguin, G.A., Rajotte, E.G., Carlo, T., 2020. Enhanced heat tolerance of viral-infected aphids leads to niche expansion and reduced interspecific competition. Nature Communications 11. http://doi.org/10.1038/s41467-020-14953-2.

Rahlff, J., Peters, J., Moyano, M., Pless, O., Claussen, C., Peck, M.A., 2017. Short-term molecular and physiological responses to heat stress in neritic copepods acartia tonsa and eurytemora affinis. Comparative Biochemistry and Physiology Part A: Molecular & Integrative Physiology 203, 348–358. http://doi.org/10.1016/j.cbpa.2016.11.001.

Roccatagliata, D., Lovrich, G.A., 1999. Infestation of the false king crab paralomis granulosa (decapoda: Lithodidae) by pseudione tuberculata (isopoda: Bopyridae) in the beagle channel, argentina. Journal of Crustacean Biology 19, 720–729.

Sasaki, M., Dam, H.G., 2021. Global patterns in copepod thermal tolerance. Journal of Plankton Research 43, 598–609. http://doi.org/10.1093/plankt/fbab044.

Schoville, S.D., Barreto, F.S., Moy, G.W., Wolff, A., Burton, R.S., 2012. Investigating the molecular basis of local adaptation to thermal stress: population differences in gene expression across the transcriptome of the copepod tigriopus californicus. BMC Evolutionary Biology 12, 170. http://doi.org/10.1186/1471-2148-12-170.

Selbach, C., Barsøe, M., Vogensen, T.K., Samsing, A.B., Mouritsen, K.N., 2020. Temperature–parasite interaction: do trematode infections protect against heat stress? International Journal for Parasitology 50, 1189–1194. http://doi.org/10.1016/j.ijpara.2020.07.006.

Sherman, E., 2008. Thermal biology of newts (notophthalmus viridescens) chronically infected with a naturally occurring pathogen. Journal of Thermal Biology 33, 27–31. http://doi.org/10.1016/j.jtherbio.2007.09.005.

Shields, J.D., 2012. The impact of pathogens on exploited populations of decapod crustaceans. Journal of Invertebrate Pathology 110, 211–224. http://doi.org/10.1016/j.jip.2012.03.011.

Somers, I., Kirkwood, G., 1991. Population ecology of the grooved tiger prawn, penaeus semisulcatus, in the north-western gulf of carpentaria, australia: growth, movement, age structure and infestation by the bopyrid parasite epipenaeon ingens. Marine and Freshwater Research 42, 349. http://doi.org/10.1071/mf9910349.

Thomas, F., Renaud, F., Guegan, J. (Eds.), 2005. Parasitism and Ecosystems. Oxford University Press.

Turner, J.T., 1981. Latitudinal patterns of calanoid and cyclopoid copepod diversity in estuarine waters of eastern north america. Journal of Biogeography 8, 369. http://doi.org/10.2307/2844757.

Uye, S.i., Murase, A., 1997. Infertility of the planktonic copepod calanus sinicus caused by parasitism by a larval epicaridian isopod. Plankton Biology & Ecology 44, 97–99.

Viechtbauer, W., 2010. Conducting meta-analyses in r with the metafor package. Journal of Statistical Software 36. http://doi.org/10.18637/jss.v036.i03.

Vinuesa, J.H., Balzi, P., 2010. Infestation of *lithodes santolla* (decapoda: Lithodidae) by *pseudione tuberculata* (isopoda: Bopyridae) in san jorge gulf, southwestern atlantic ocean. Marine Biology Research 6, 608–612. http://doi.org/10.1080/17451000903478327.

Wertheim, B., 2022. Adaptations and counter-adaptations in drosophila host–parasitoid interactions: advances in the molecular mechanisms. Current Opinion in Insect Science 51, 100896. http://doi.org/10.1016/j.cois.2022.100896.

Wertheim, B., Kraaijeveld, A.R., Schuster, E., Blanc, E., Hopkins, M., Pletcher, S.D., Strand, M.R., Partridge, L., Godfray, H.C.J., 2005. Genome Biology 6, R94. http://doi.org/10.1186/gb-2005-6-11-r94.

Willett, C.S., 2010. Potential fitness trade-offs for thermal tolerance in the intertidal copepod tigriopus californicus. Evolution 64, 2521–2534. http://doi.org/10.1111/j.1558-5646.2010.01008.x.

Williams, J.D., Escalante, M., Shanks, A.L., 2022. Identification and observations of parasitic isopod larvae (isopoda: Epicaridea) from the northeastern pacific: pelagic distribution and association with copepod intermediate hosts. Journal of Crustacean Biology 42, ruac045. http://doi.org/10.1093/jcbiol/ruac045.

Yang, C., Shan, B., Liu, Y., Wang, L., Liu, M., Yao, T., Sun, D., 2022. Transcriptomic analysis of male three-spot swimming crab (portunus sanguinolentus) infected with the parasitic barnacle diplothylacus sinensis. Fish & Shellfish Immunology 128, 260–268. http://doi.org/10.1016/j.fsi.2022.07.074.

Yu, C., Xu, W., Li, X., Jin, J., Zhao, X., Wang, S., Zhang, Z., Wei, Y., Chen, Q., Li, Y., 2021. Comparative transcriptome analysis of chinese grass shrimp (palaemonetes sinensis) hepatopancreas under ectoparasitic isopod (tachaea chinensis) infection. Fish & Shellfish Immunology 117, 211–219. http://doi.org/10.1016/j.fsi.2021.07.018.

